# Transcriptomic profiling and microsatellite identification in cobia (Rachycentron canadum), using high throughput RNA-sequencing

**DOI:** 10.1101/2020.12.04.374918

**Authors:** David Aciole Barbosa, Bruno C. Araújo, Giovana Souza Branco, Alexandre S. Simeone, Alexandre W. S. Hilsdorf, Daniela L. Jabes, Luiz R. Nunes, Renata G. Moreira, Fabiano B. Menegidio

## Abstract

Cobia (*Rachycentron canadum*) is a marine teleost species with great productive potential worldwide. However, the genomic information currently available for this species in public databases is limited. Such lack of information hinders gene expression assessments that might bring forward novel insights into the physiology, ecology, evolution, and genetics of this potential aquaculture species. In this study, we report the first *de novo* transcriptome assembly of *R. canadum* liver, improving the availability of novel gene sequences for this species. Illumina sequencing of liver transcripts generated 1,761,965,794 raw reads, which were filtered into 1,652,319,304 high-quality reads. *De novo* assembly resulted in 101,789 unigenes and 163,096 isoforms, with an average length of 950.61 and 1,617.34 nt, respectively. Moreover, we found that 126,013 of these transcripts bear potentially coding sequences, and 125,993 of these elements (77.3%) correspond to functionally annotated genes found in six different databases. We also identified 701 putative ncRNA and 35,414 putative lncRNA. Interestingly, homologues for 410 of these putative lncRNAs have already been observed in previous analyzes with *Danio rerio*, *Lates calcarifer*, *Seriola lalandi dorsalis*, *Seriola dumerili* or *Echeneis naucrates*. Finally, we identified 7,894 microsatellites related to cobia’s putative lncRNAs. Thus, the information derived from the transcriptome assembly described herein will likely assist future nutrigenomics and breeding programs involving this important fish farming species.

## 1. INTRODUCTION

Currently, aquaculture is the most prominent food production industry, with significant growth worldwide. Global aquaculture production increased by 5.3% percent per year between 2001-2018, with a historical record of 114.5 million tons of farmed species, including almost 17.7 million tons due to finfish production (FAO 2020). Marine aquaculture plays an essential role in the effort of providing the increasing world’s demand for animal-based protein. Cobia (*Rachycentron canadum*) is a carnivorous marine fish of worldwide distribution, and it is the sole representative of the Rachycentridae family, among farmed fish species. Currently, *R. canadum* is regarded as the most promising marine fish species in Brazil, mostly due to its fast growth rate (reaching about 4 to 6 kg per year), excellent meat quality (with regards to color, texture and flavor), and high market value (Arnold et al. 2002; Benetti et al. 2008; Nunes 2014). However, cobia production is still hindered by the lack of nutrition information, which constraints this species’ productivity in industrial aquaculture operations (Fraser and Davies 2009).

Next-generation sequencing (NGS) studies have become an essential molecular tool in aquaculture, assisting in the production of several commercial fish species, such as *Sparus aurata* (Calduch-Giner et al. 2013), *Dicentrarchus labrax* (Magnanou et al. 2014), and *Salmo salar* (Glencross et al. 2015; Andrew et al. 2021). *De novo* transcriptome assembly can be used in several different contexts, like genomics/gene expression analyses and may be applied in many key areas of study, such as conservation genetics, selective breeding, reproductive biology, and nutrition (Leaver et al. 2008; Calduch-Giner et al. 2013; Fox et al. 2014). For example, identifying genes associated with proteins and lipid metabolism in the liver can assist in the development of specific diets to improve the productive chain of commercial aquaculture species, such as cobia. Unfortunately, genomic information in cobia is still scarce, limiting the development of such studies. Thus, to help overcome such limitations, the present manuscript describes an assembled/annotated reference transcriptome of hepatic cells in cobia juveniles. The information contained in this dataset contributes to improve our knowledge regarding the biological and physiological aspects of this fish species and establishes a solid foundation for future studies involving population genomics, breeding programs, and nutrigenomics involving this important marine farming fish.

## 2. MATERIALS AND METHODS

### 2.1. Sample collection and RNA preparation

Ninety cobia juveniles (128.85 ± 18.43 g) were obtained from a commercial hatchery (Redemar Alevinos, SP, Brazil) and randomly allocated in three 2,000 L tanks. The animals were kept under a mean temperature of 23±1.5 °C and a photoperiod of 12L:12D throughout the trial, at the Marine Biology Center of the University of São Paulo (CEBIMar). Animals were equally hand-fed twice a day, until apparent satiety, with a commercial marine fish diet (Guabipirá, Guabi Nutrição e Saúde Animal S.A., SP, Brazil). After six weeks (42 days), fish were anesthetized with benzocaine (0.4 g * mL^-1^) and then euthanized by spinal cord section. Hepatic tissue samples from all experimental animals were collected, immediately frozen in liquid nitrogen, and subsequently stored at −80 °C for further analyses. Total RNA from hepatic tissue samples was extracted using Rneasy Lipid Tissue kit (Qiagen), following the manufacturer’s instructions. RNA samples had their concentration determined with a NanodropTM Spectrophotometer (ND-1000). RNA integrity was assessed using a 2100 Bioanalyzer System (Agilent Technologies, USA). This study’s experimental procedures were conducted according to the guidelines and approval of the Institutional Animal Care and Use Ethics Committee (#008/2017).

### 2.2. Library construction and sequencing

RNA extracted from liver samples from all animals (90 fish, 30 per tank), were equally diluted to 1,000 ng/μl concentration and pooled for library construction, using the TruSeq RNA Sample Preparation kit, according to the manufacturer’s specifications (Illumina Inc., USA). Library quality was validated using a Bioanalyzer 2100 (Agilent Technologies, USA), and only samples with an RNA Integrity Number (RIN) equal or above 7.5 were used. Finally, paired-end sequencing (2×75 bp) of these cDNA libraries was conducted in an Illumina Nextseq® platform, according to the manufacturer’s recommendations. Minimum information about any (x) sequence (MIxS) data for this study is available in Supplementary Table ST1-S1.

### 2.3. Bioinformatics Analysis of Raw Data

FASTQ format raw sequencing data was processed in a Public Galaxy Server available at https://usegalaxy.eu. Initially, the quality of raw sequences was assessed using FastQC (Andrews 2010) and MultiQC (Ewels et al. 2016). Fastp (Chen et al. 2018) was then used to remove low-quality reads (Q<30), adapters, and other contaminant sequences. Trinity software (Haas et al. 2013) was then used for *de novo* transcriptome assembly of filtered reads, and assembly metrics were obtained using the TrinityStats script. Transcriptome completeness was finally assessed using the Benchmarking Universal Single-Copy Orthologs (BUSCO) software (Seppey et al. 2019) based on OrthologDB version 9 (Zdobnov et al. 2017).

### 2.4. Transcriptome Annotation

Elements of the assembled transcriptome were functionally annotated by similarity searches (blastn, e-value ≤ e-5) performed against RefSeq RNA and “*Lates/Seriola*” - a custom database containing RefSeq transcripts from the cobia-related species *Lates calcarifer*, *Seriola dumerili*, and *Seriola lalandi dorsalis*, which are in the Kegg’s Organisms Complete Genomes (see https://www.genome.jp/kegg/catalog/org_list.html). Additional annotations were obtained with the aid of the Eukaryotic Non-Model Transcriptome Annotation Pipeline (EnTAP) (Hart et al. 2020). Contigs were queried (blastx; using e-value ≤ e-5 and ≥ 50% coverage) for similarity against the National Center for Biotechnology Information non-redundant protein database (NCBI nr), NCBI proteins reference database (RefSeq), the curated Swiss-Prot database from UniProt Knowledgebase (UniProtKB) (UniProt Consortium 2019), and the EggNOG proteins database (Huerta-Cepas et al. 2016). The EggNOG hits also helped to assign biological function to individual elements, identifying their respective Gene Ontology (GO) (The Gene Ontology Consortium 2019) and KEGG (Kyoto Encyclopedia of Genes and Genomes) (Kanehisa and Goto 2000; Kanehisa 2019; Kanehisa et al. 2021) terms. The EnTAP functional annotation process was carried out using a Dugong container environment (Menegidio et al. 2018). Transcripts not annotated by EnTAP were evaluated using the cmscan program (default parameters) by Infernal (Nawrocki and Eddy 2013), for classification in the different families of non-coding RNAs, defined in the Rfam database (Kalvari et al. 2018).

### 2.5. Coding Potential Calculator and lncRNA Discovery

Transcripts not annotated by EnTAP and Infernal were evaluated for their respective coding potential (CP) with the aid of three tools: Coding Potential Calculator (CPC2) (Kang et al. 2017), Coding-Potential Assessment Tool (CPAT) (Wang et al. 2013) and RNASamba (Camargo et al. 2020). The transcripts identified as having non-coding potential by all of these tools were separated for functional annotation analysis. In this work, we considered putative lncRNAs transcripts with ≥ 200 nt that were identified as non-coding by all of CP tools and not annotated by EnTAP / Infernal. To discover conserved interspecies lncRNAs, we aligned putative lncRNA sequences against the Zebrafish lncRNA Database (ZFLNC; Hu et al. 2018) using blastn (Boratyn et al. 2013) with a cut-off value ≤ e-5 and ≥50% identity (Fan et al.2018). Similar, blastn searches were employed against ncRNA sequences available at Ensembl from *Danio rerio*, *Lates calcarifer*, *Echeneis naucrates*, *Seriola lalandi dorsalis* and *S. dumerili*.

### 2.6. Detection of SSRs in lncRNAs

The MIcroSAtellite (MISA) software (Beier et al. 2017) was used to identify microsatellites in the putative lncRNAs sequences. The Simple Sequence Repeats (SSR) loci detection was done by searching for two-to six-nucleotide motifs, with a minimum of 1/10, 2/6, 3/5, 4/5, 5/4 and 6/4 (motifs/repeats), as suggested by Gui et al., (2013).

## 3. RESULTS AND DISCUSSION

### 3.1 Transcriptome assembly and completeness

Sequencing of the cDNA libraries derived from *R. canadum* liver material resulted in 1,761,965,794 raw reads. After high-quality-read selection and trimming, we were left with a total of 1,652,319,304 reads (93.77% of raw reads), which were used for *de novo* transcriptome assembly, using Trinity software (Haas et al. 2013). General features of the *R. canadum* liver transcriptome are summarized in Table 1, consisting of 101,789 unigenes and 163,096 isoforms (likely derived from cryptic transcription start sites, alternative splicing or differential polyadenylation events). The median (N50)/average length of these elements was 7,843/1,617.34 nt for unigenes and 2,312/950.61 nt for isoforms. A total of 95,075 transcripts (58.29%) were ≥500 nt. Identification of 83.8% of the complete universal genes (3,839 out of the total 4,584 genes from Actinopterygii odb9 lineage) supported the high quality and completeness of this transcriptome assembly (Fig. 1a). Among the 3,839 conserved BUSCO genes, 38.1% were single copy, while 45.7% were duplicated (Supplementary Table ST1-S2).

**Fig. 1.**
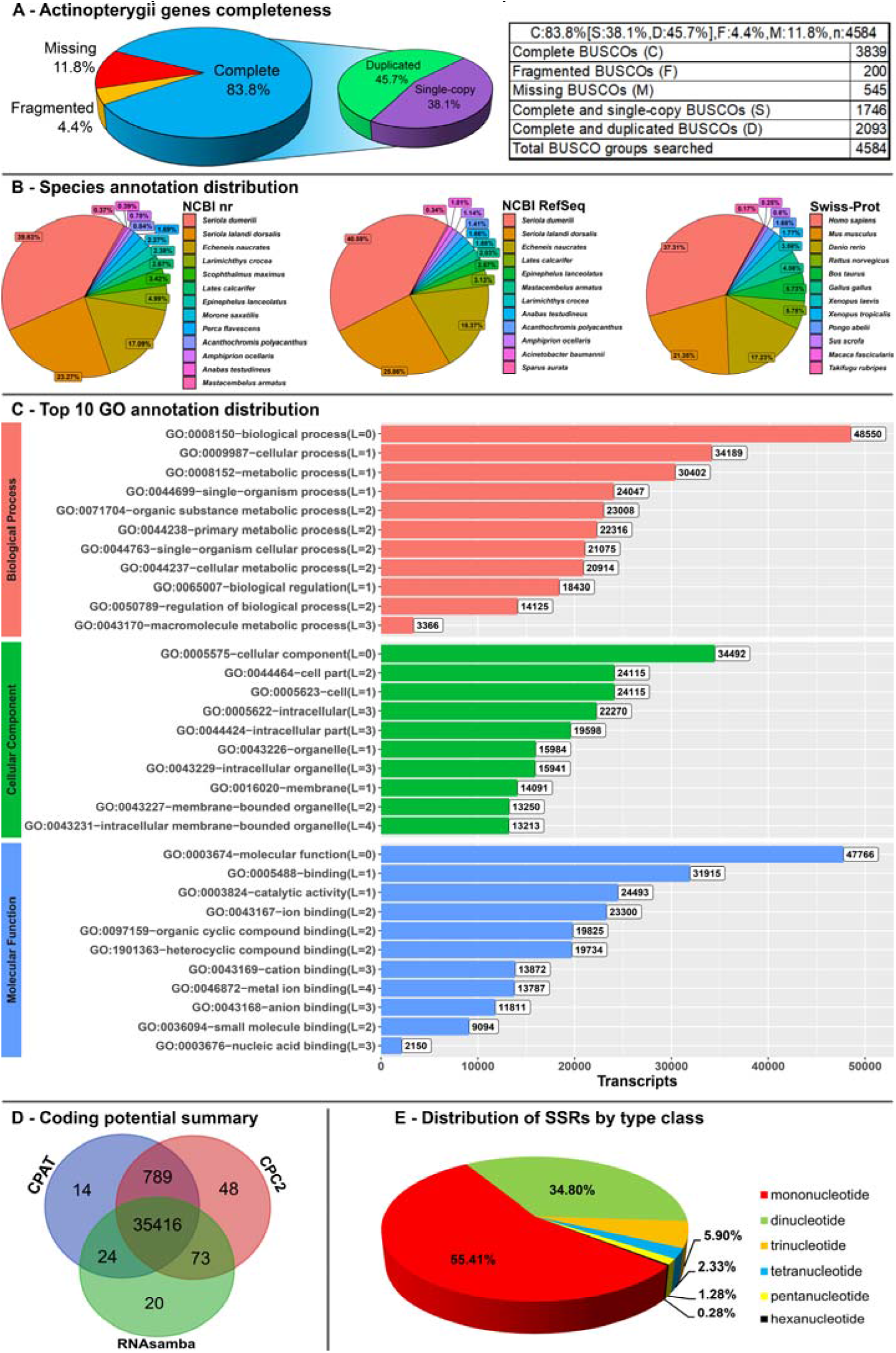
Completeness and homology search of *Rachycentron canadum* liver transcripts. (A) Percentage of completeness on the core set of genes from *R. canadum* based on Actinopterygii database (orthodb9), using BUSCO. (B) Species annotation distribution for the best hits from NCBI nr, NCBI RefSeq and Swiss-Prot databases. (C) Gene ontology distribution for Biological Process, Cellular Component, Molecular Function categories of assembled transcripts from the *R. canadum* liver transcriptome. (D) Distribution of transcripts classified as non-protein coding elements. (E) Distribution of microsatellites (SRRs) identified in the putative lncRNA sequences, based on their respective classes.

**Table 1.** Summary of the *de novo* transcriptome assembly for *Rachycentron canadum*.

| Illumina sequencing | Raw | High-Quality |
| --- | --- | --- |
| Number of reads | 1,761,965,794 | 1,652,319,304 |
| Assembly | Unigenes | Isoforms |
| Total sequences | 101,789 | 163,096 |
| N50 | 7,843 | 2,312 |
| Average contig length (bp) | 1,617.34 | 950.61 |
| Median contig length (bp) | 704 | 391 |
| Total assembled bases | 263,781,171 | 96,761,286 |
| CG% | 44.5 |  |
| Coding potential | Transcripts |  |
| Undetermined transcripts | 968 |  |
| Coding transcripts | 126,013 |  |
| Non-coding transcripts | 701 |  |
| Putative long non-coding transcripts | 35,414 |  |
| Functional Annotation | Annotated Sequences |  |
| NCBI nr | 29,155 |  |
| NCBI RefSeq RNA | 122,741 |  |
| Swiss-Prot | 15,507 |  |
| Lates/Seriola | 118,570 |  |
| GO-Biological Process | 48,550 |  |
| GO-Molecular Function | 47,766 |  |
| GO-Cellular Component | 34,492 |  |
| Kegg | 23,936 |  |
| Infernal | 699 |  |
| ZFLNC | 206 |  |
| ncRNA Ensembl DB | 257 |  |

### 3.2 Functional annotation

Transcriptome annotation against a series of databases (NCBI nr, NCBI RefSeq RNA, Swiss-Prot, GO, KEGG, *Lates/Seriola*) resulted in functional assignment for 125,993 transcripts (77.3%). Most sequence homologies were found against NCBI RefSeq RNA (122,741 transcripts, or 97.4%), followed by *Lates* and *Seriola* species databases (118,570 transcripts, or 94.1%), NCBI nr (29,155 transcripts, or 23.14%), NCBI RefSeq (25,728, or 20.42%) and Swiss-Prot (15,507 transcripts, or 12.30%) (Table1; Supplementary Table ST1-S3a-S5). Sequence homologies identified by EnTAP were distributed through many bony fish species, of which *Seriola dumerili* was the most frequent (nr = 39.83%, RefSeq = 40.59%), followed by *Seriola lalandi dorsalis* (nr = 23.27%, RefSeq = 25.86%), *Echeneis naucrates* (nr 17.9%, RefSeq = 18.37%) and *Larimichthys crocea* (nr = 4.99%, RefSeq = 3.13%) (Fig. 1b). Most transcripts (75,060) were functionally annotated with GO terms by eggNOG (Huerta-Cepas et al. 2016), which assigned 48,550 transcripts (65%) to biological processes, 34,492 to cellular components (46%) and 47,766 to molecular functions (64%). The ten most representative functional groups within each category are shown in Fig. 1c. A total of 23,936 isoforms were annotated into at least one KEGG pathway term (Table1; Supplementary Table ST1-S6).

The final functional annotation allowed us to identify several marker genes for future nutrigenomics initiatives, including elements involved in the following GO Biological Processes: (i) GO:0007586 - Digestion of Nutrients (ex.: Carboxypeptidases, Trypsin and Trypsin-like homologues, Chymotrypsin-like Elastase Family members, etc.); (ii) GO:0042445 - Proteins Involved with Hormone Function/Metabolism (ex. Calcitonin, Estrogen Receptors, Hepcidins, Insulin/Glucagon homologues and receptors, etc.); (iii) GO:0006629/GO:0005975 Lipid/Carbohydrate Metabolic Processes (ex. Phospholipases A/B/C/D, Bile Salt-stimulated Lipases, Chitinases, Galactosidases, Alpha/Beta Glucosidases, etc.); (iv) GO:0044765 - Nutrient Transport (ABC Transporters, Amino Acid Permeases, Apolipoprotein/Hemoglobin homologues, etc.); (v) GO:0048644 - Muscle Structure and Morphogenesis (ex. Actin/Myosin/Formin/Growth Factor homologues, etc,) and (vi) GO:0048565 - Digestive Tract Development (ex. Kruppel-like Factors, Digestive Organ Expansion Factor homologues, Pancreatic and Duodenal Homeobox 1-containing proteins, Hepatocyte Growth Factors, etc), among others (see Supplementary Table ST1-S3b, for details).

### 3.3 Known Non-Coding RNAs

For the 37,103 transcripts not annotated in the previous steps, the Infernal tool suite (Nawrocki and 2013) was used to filter the presence of known non-coding RNAs available from the RFAM database (Kalvari et al. 2018). The cmscan script allowed annotation of 33,677 such sequences. Among these, we obtained 699 significant hits (based on an e-value threshold of 0.001), allowing their classification as putative non-coding RNAs (Table1; Supplementary Table ST1-S7). Among the ncRNAs identified in our *R. canadum* transcriptome are small nucleolar RNAs (snoRNAs), such as SNORD5, SNORD21, SNORD27, SNORD31, SNORD36, SNORD48, SNORD52, SNORD63, SNORD78, SNORD88 and SNORD103 - which have also been described in *Danio rerio*, *Gasterosteus aculeatus*, *Tetraodon nigroviridis* and *Oryzias latipes*, according to the snoOPY database (http://snoopy.med.miyazaki-u.ac.jp). Cajal small body RNAs (scaRNAs), which are involved in the modification of snoRNAs, were also identified in our data. Among the scaRNAs, we found a homologue for SCARNA7, a C/D RNA box, which localizes to Cajal bodies in HeLa cells and is conserved in all vertebrates (Marz et al. 2011) MicroRNA (miRNA), accounted for 170 non-coding transcripts, distributed across 111 miRNA types. MicroRNAs have been found to be related to several relevant biological aspects in fish, including regulation of growth, muscle, vascular and cardiac tissues, among many others (Rasal et al. 2016). Moreover, Herkenhoff et al. (2018) suggested that miRNAs may also serve as biomarkers for selection of adaptive traits for aquaculture.

### 3.4 Novel Non-Coding RNAs and Long Non-Coding RNAs

Next, transcripts not annotated by EnTAP and Infernal were subjected to a coding potential (CP) analysis, using four different CP calculator tools: Coding Potential Calculator (CPC2) (Kang et al. 2017), Coding-Potential Assessment Tool (CPAT) (Wang et al. 2013) and RNASamba (Camargo et al. 2020), which subdivided these novel elements into coding and non-coding transcripts (Supplementary Table ST1-S8). Among the remaining 36,404 unannotated elements, 0.054% were identified as protein coding (representing 20 putative new genes), 2.65% were of undetermined nature (968 transcripts) and 97.29% were classified as non-protein coding elements (35,416 transcripts). The distribution of transcripts classified as non-protein coding elements (non-coding RNAs) can be observed in Fig. 1d (the 35,416 elements specified at this point are represented by the intersection of all CP tools). Among these, 35,414 were larger than 200 bp, and were finally classified as putative long-noncoding RNAs (lncRNAs) (Supplementary Table ST1-S9). Most cobia lncRNAs (26,255, 74.14%) are larger than 400 bp; 5,582 of them are ≥ 600 bp (15.76%), 1,871 (5.28%) are ≥800 bp and 778 (2.20%) are ≥1,000 bp, while 928 (2.62%) have sizes ≥ 2,000 bp.

### 3.5 Long Non-Coding RNAs Annotation

We performed an orthologous analysis of our putative lncRNAs using the ZFLNC database and a custom ncRNA sequences database from *Danio rerio*, *Lates calcarifer*, *Seriola lalandi dorsalis*, *Seriola dumerili*, *Echeneis naucrates* available at Ensembl (see Methods, for details). Among the 206 ZFLNC hits, the most common annotations were ZFLNCT01535, ZFLNCT11671, ZFLNCT19022 (matches with 5 transcripts, each), ZFLNCT02442 (matches with 4 transcripts), ZFLNCT13035 and ZFLNCT16489 (each matching three putative lncRNAs). Interestingly, among the 79 hits for *D. rerio* ncRNA sequences present in Ensembl, 57 are annotated as long intervening noncoding RNAs/Long intergenic noncoding RNAs (lincRNAs). The remaining hits correspond to antisense (5), retained_intron (2), misc_RNA (1), sense intronic (1) snoRNA (1) and 12 of them as processed transcript (Supplementary Table ST1-S9). As mentioned above, the majority of transcripts described herein displayed significant similarity to coding DNA of fish species phylogenetically related to cobia, such as *Seriola dumerili*, *Seriola lalandi dorsalis*, *Echeneis naucrates* and *Lates calcarifer*, during our initial annotation efforts. The lncRNA conservation search performed here showed that 157 of 159 hits obtained from *E. naucrates* ncRNAs correspond to sequences already identified as lncRNA in this species (the other 2 hits are annotated as small nucleolar RNAs U85 and SNORD10). On the other hand, only few hits were found from *Seriola* spp. and *L. calcarifer*: (i) one snoRNA appeared only for *L. calcarifer* and is the same lncRNA transcript matching *E. naucrates* SNORD10; (ii) one misc_RNA, called 7SK RNA, was found for the same transcript for these organisms – which is also the same hit found from *D. rerio* (Supplementary Table ST1-S9).

### 3.6 lncRNA Microsatellites

Sequences from all 35,414 putative lncRNA transcripts were used to discover potential microsatellites in the cobia genome with the aid of MISA. The SSR loci detection was performed by searching for two to six nucleotide motifs, with a minimum of 6,5,5,4 and 4 repeats, respectively. A total of 7,894 microsatellites were detected in the putative lncRNAs (Supplementary Table ST1-S10). Among the microsatellites, mono-nucleotide motifs were the most abundant type detected in lncRNAs (55.41%). Other motifs included di-nucleotide (34.80%), tri-nucleotide (5.9%), tetra-nucleotide (2.33%), penta-nucleotide (1.28%) and hexa-nucleotide (0.28%) motifs (Fig. 1e). The mono-nucleotide repeat T was the most abundant motif detected (50.07%), followed by A (46.27%), C (2.29%) and, finally, G (1.37%) (Supplementary Table ST1-S10).

## 4. CONCLUSION

Our study has built the first liver transcriptome assembly of this important commercial species, providing an important tool for further research with cobia. The availability and deposition of the transcriptome sequence allows to access novel gene sequences, contributing to gene expression assessments, and consequently improving the knowledge regarding cobia physiology and nutrition, since the liver can be considered the main lipogenic tissue in fish. In addition, the provided assembly and genetic markers dataset will be essential as a base for future nutrigenomics projects involving, genetic breeding programs and marker-assisted selection for this species.

## DECLARATIONS

### Availability of data and material

Sequencing raw data were deposited in the Sequence Read Archive (SRA) repository of the National Center for Biotechnology Information (NCBI), under accession number SRR13009897, SRR13009896, SRR13009895, SRR13009894, SRR13009893, SRR13009892, SRR13009891, SRR13009890, SRR13009889, SRR13009888, SRR13009887 and SRR13009886, associated to the BioProject numbers PRJNA675281 and BioSamples numbers SAMN16708758, SAMN16708759, SAMN16708760, SAMN16708761, SAMN16708762, SAMN16708763, SAMN16708764, SAMN16708765, SAMN16708766, SAMN16708767, SAMN16708768 and SAMN16708769. The Transcriptome Shotgun Assembly (TSA) project has been deposited at DDBJ/EMBL/GenBank under accession number GIWT00000000. The version described in this paper is the first version, GIWT00000000.1. Supplementary Table S1 is available from the Figshare repository (10.6084/m9.figshare.14522781.v2). Additional data derived from this study (including all intermediate data) are also available from the Open Science Framework (OSF) repository (DOI: 10.17605/OSF.IO/BV3WA). Details about the softwares and databases used are available in Supplementary Table ST1-S11.

### Authors’ contributions

B.C.A. sampled the specimens. B.C.A., G.S.B., A.W.S.H. and R.G.M. performed molecular analyses and sequencing. D.A.B., A.S.S., D.L.J., L.R.N. and F.B.M. assembled and evaluated the transcriptome assembly and annotation. All authors wrote the paper. All authors read and approved the final version of the manuscript.

### Ethics approval

This study’s experimental procedures were conducted according to the guidelines and approval of the Mogi das Cruzes University Institutional Animal Care and Use Ethics Committee (#008/2017).

